# A full-length transcriptome dataset of normal and *Nosema ceranae*-challenged midgut tissues of eastern honeybee workers

**DOI:** 10.1101/2020.03.18.997981

**Authors:** Yu Du, Yuanchan Fan, Huazhi Chen, Jie Wang, Cuiling Xiong, Yanzhen Zheng, Dafu Chen, Rui Guo

## Abstract

*Apis cerana cerana* is a subspecies of eastern honeybee, *Apis cerana*, and it plays a vital role in ecological maintenance in China. However, *A. c. cerana* is threatened by many pathogenic microorganisms including *Nosema ceranae*, a widespread fungal parasite that infected worldwide colonies. In this article, un-challenged (AcCK1, AcCK2) and *N. ceranae*-challenged midguts of *A. c. cerana* workers (AcT1, AcT2) were sequenced utilizing Nanopore long-read sequencing technology. Totally, 11,727,628, 6,996,395, 14,383,735 and 11,580,154 raw reads were yielded from AcCK1, AcCK2, AcT1 and AcT2; the average lengths were 1147 bp, 908 bp, 992 bp and 1077 bp, and the average N50 were 1308 bp, 911 bp, 1079 bp and 1192 bp. The length distribution of was ranged 1 kb to more than 10 kb. Additionally, the quality (Q) score distribution of raw reads was among Q7~Q17. Further, 11,617,144, 6,940,895, 14,277,240 and 11,501,562 clean reads were respectively obtained from AcCK1, AcCK2, AcT1 and AcT2, and among them 78.40%, 82.50%, 79.05% and 80.20% were identified as full-length clean reads. In addition, full-length clean reads from AcCK1, AcT1, AcT2 and AcCK2 were ranged from 1 kb to more than 10 kb in length. Finally, the length distribution of redundant reads-removed full-length transcripts was among 1 kb~5 kb.

**Value of the data:**

♦ This dataset enables better understanding the complexity of *A. c. cerana* transcriptome.
♦ Current dataset contributes to identification of genes and transcripts engaged in response of eastern honeybee to *N. ceranae* stress.
♦ The data provides a valuable genetic resource for deciphering alternative splicing and polyadenylation of *A. c. cerana* mRNAs involved in host response to *N. ceranae* challenge.
♦ The reported data is beneficial for uncovering the molecular mechanism regulating interaction between eastern honeybee and microsporidian.

## 1. Data Description

The shared full-length transcriptome data were generated from un-challenged and *N. ceranae*-challenged midgut tissues of *A. c. cerana* workers. Totally, 11,727,628, 6,996,395, 14,383,735 and 11,580,154 raw reads were respectively yielded from AcCK1, AcCK2, AcT1 and AcT2 (**Table 1**); the average lengths were 1147 bp, 908 bp, 992 bp and 1077 bp, and the average N50 were 1308 bp, 911 bp, 1079 bp and 1192 bp (**Table 1**). Raw reads of AcCK2 were ranged from 1 kb to 10 kb in length, with the most abundant length of 1 kb (**Figure 1B**), while the length of raw reads from AcCK1, AcT1 and AcT2 was ranged from 1 kb to more than 10 kb, and the most abundant length was also 1 kb (**Figure 1A,C,D**). In addition, as **Figure 2** shown, the quality (Q) score distribution of raw reads from the aforementioned four groups was among Q7~Q17, with the highest percentage of Q11. From AcCK1, AcCK2, AcT1 and AcT2, 11,617,144, 6,940,895, 14,277,240 and 11,501,562 clean reads were respectively obtained (**Table 2**); among them 78.40%, 82.50%, 79.05% and 80.20% were identified as full-length clean reads (**Table 2**). Additionally, full-length clean reads from AcCK1, AcT1, AcT2 and AcCK2 were ranged from 1 kb to more than 10 kb in length, with the largest group of 1 kb (**Figure 3**). Moreover, as **Figure 4** presented, after removing redundant reads, the length of remaining transcripts was distributed among 1 kb~5 kb, with the highest percentage of 1 kb for AcCK2 and AcT1 and 2 kb for AcCK1 and AcT2.

**Table 1.** An overview of raw data derived from Nanopore long-read sequencing

| Sample | Reads number | Base number | N50<br>(bp) | Mean<br>length<br>(bp) | Maximum length<br>(bp) |
| --- | --- | --- | --- | --- | --- |
| AcCK1 | 11,727,628 | 13,456,485,261 | 1308 | 1147 | 16,647 |
| AcCK2 | 6,996,395 | 6,356,440,568 | 911 | 908 | 9864 |
| AcT1 | 14,383,735 | 14,272,926,859 | 1079 | 992 | 65,767 |
| AcT2 | 11,580,154 | 12,480,305,079 | 1192 | 1077 | 13,293 |

**Table 2.** A summary of full-length clean reads

| Sample | Number of<br>clean reads | Number of full-length<br>clean reads | Percentage of full-length<br>clean reads |
| --- | --- | --- | --- |
| AcCK1 | 11,617,144 | 9,108,082 | 78.40% |
| AcCK2 | 6,940,895 | 5,726,448 | 82.50% |
| AcT1 | 14,277,240 | 11,286,184 | 79.05% |
| AcT2 | 11,501,562 | 9,224,291 | 80.20% |

**Figure 1.**
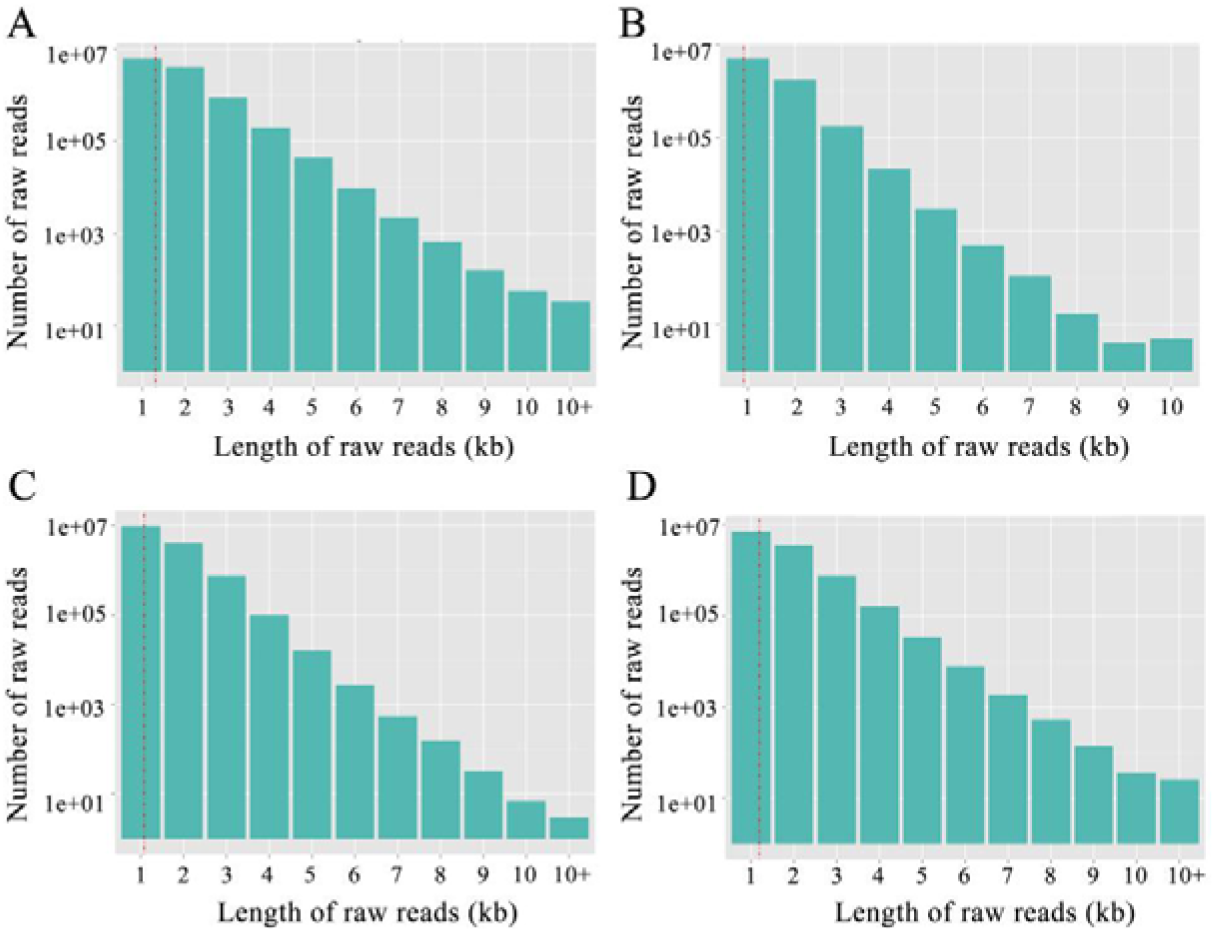
Length distribution of raw reads generated from Nanopore long-read sequencing. (A) Raw reads from AcCK1; (B) Raw reads from AcCK2; (C) Raw reads from AcT1; (D) Raw reads from AcT2.

**Figure 2.**
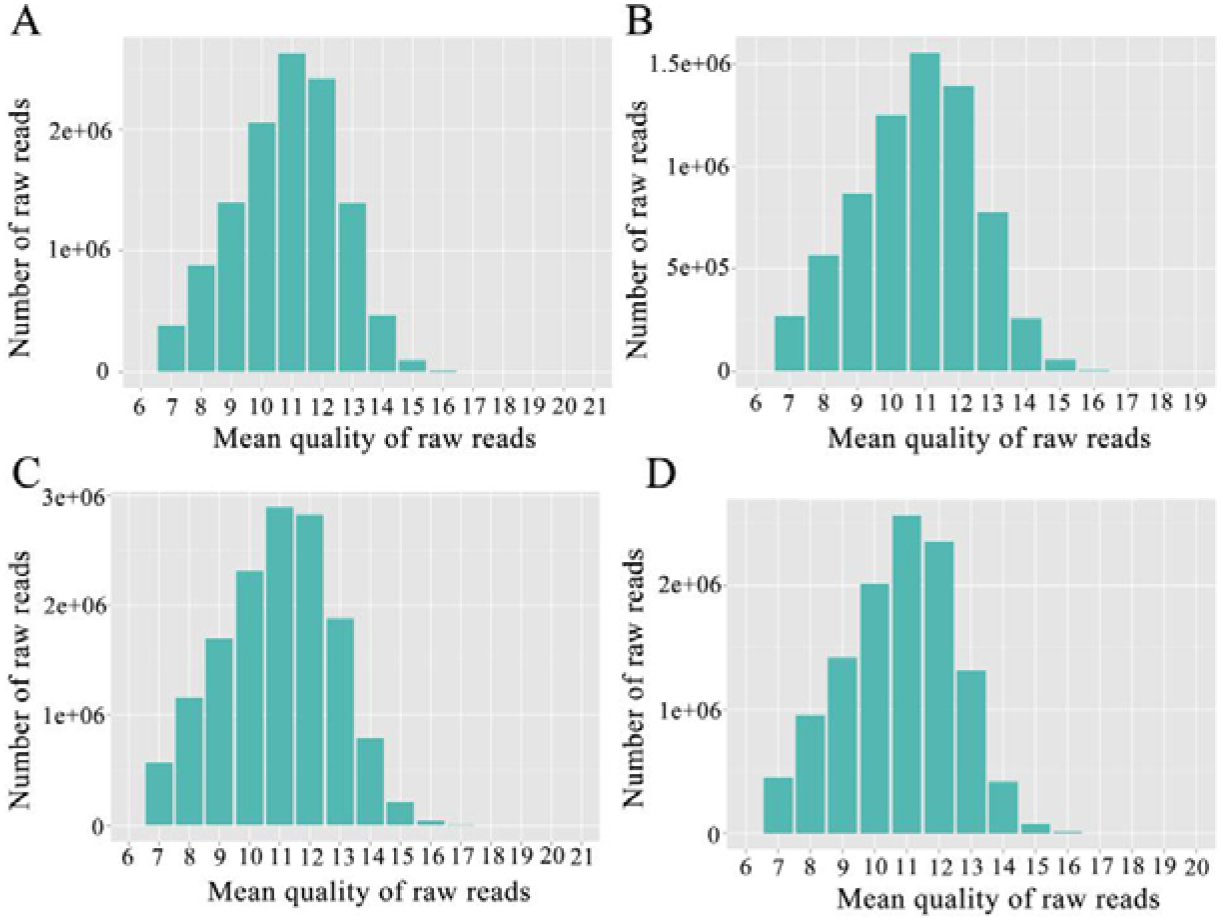
Quality distribution of raw reads derived from Nanopore long-read sequencing. (A) Raw reads from AcCK1; (B) Raw reads from AcCK2; (C) Raw reads from AcT1; (D) Raw reads from AcT2.

**Figure 3.**
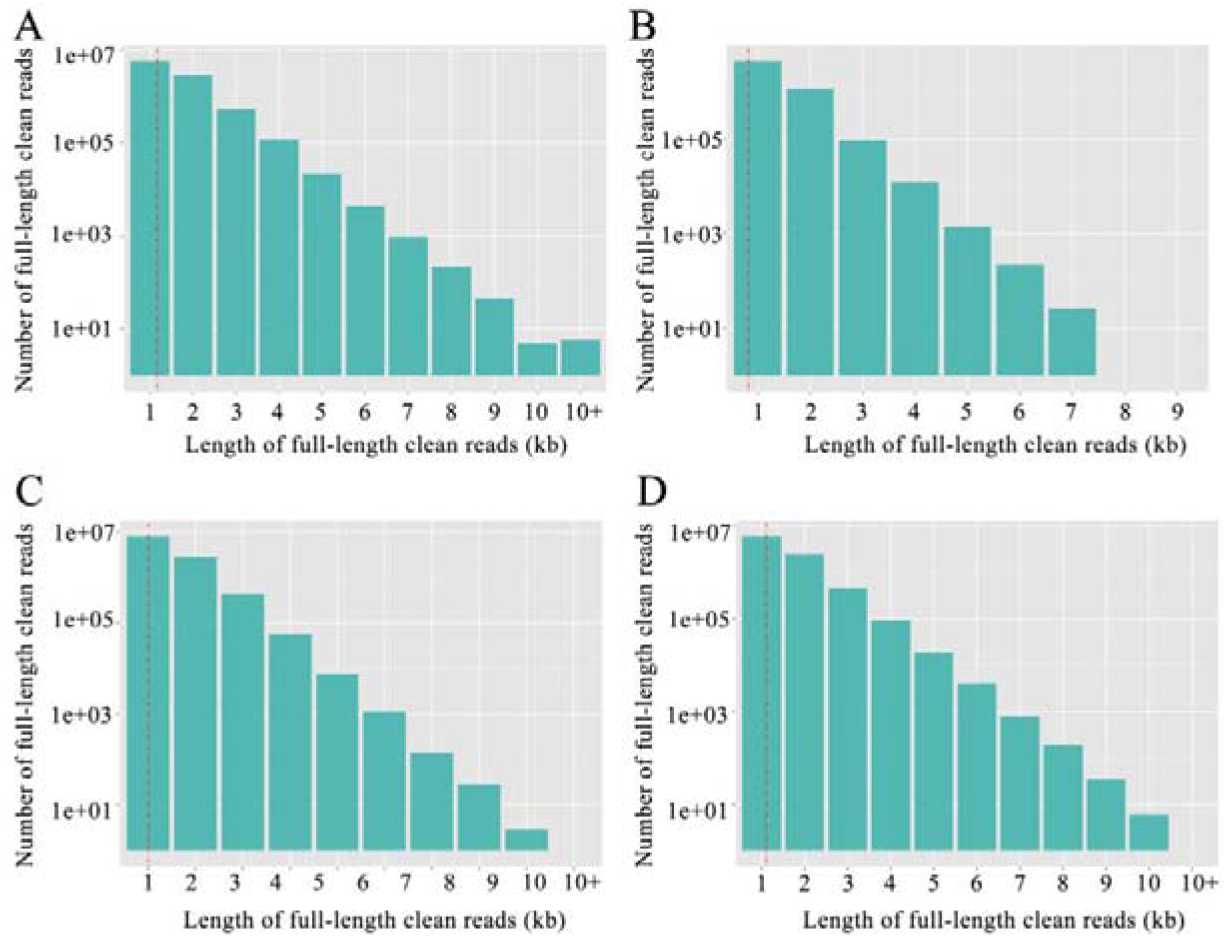
Length distribution of full-length clean reads. (A) Full-length clean reads from AcCK1; (B) Full-length clean reads from AcCK2; (C) Full-length clean reads from AcT1; (D) Full-length clean reads from AcT2.

**Figure 4.**
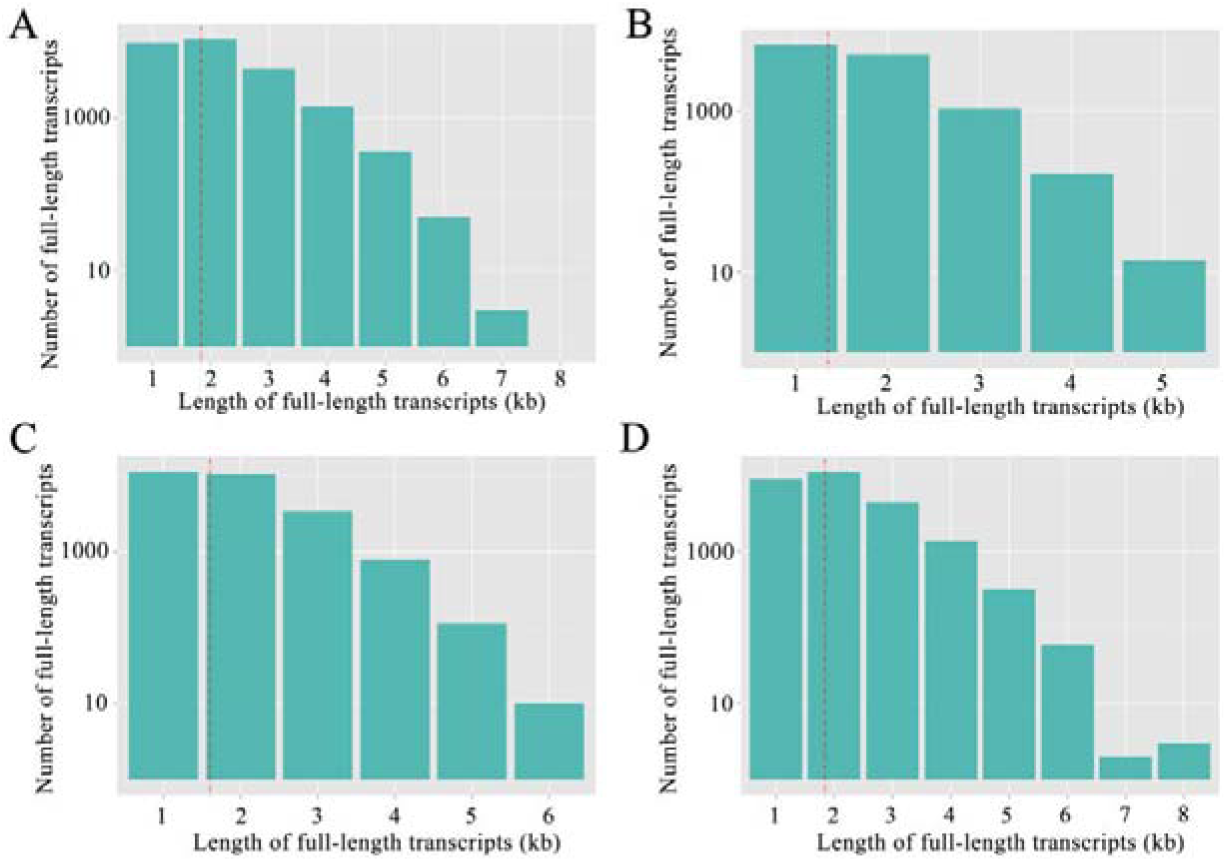
Length distribution of full-length transcripts after removing redundant reads. (A) Redundant reads-removed full-length transcripts from AcCK1; (B) Redundant reads-removed full-length transcripts from AcCK2; (C) Redundant reads-removed full-length transcripts from AcT1; (D) Redundant reads-removed full-length transcripts from AcT2.

## 2. Experimental Design, Materials, and Methods

### 2.1. Experimental design and sample collection

*N. ceranae* spores were previously purified following the method described by Cornman et al. [1] with some minor modifcations [2]. A small quantity of spores was examined by PCR detection and verified to be mono-specific [3] using previously developed specific primers [4]. The purified spores were immediately used for artificial inoculation of workers in treatment groups. Artificial inoculation of bees and preparation of midgut samples were performed following our previously established method [3,5]. (1) Frames of sealed brood gained from a healthy colony of *A. c. cerana* located in the teaching apiary of College of Animal Sciences (College of Bee Science), Fujian Agriculture and Forestry University were raised in an incubator at 34±0.5 °C, 50% RH to provide newly emerged *Nosema*-free workers; the emergent workers were carefully transferred to cages in groups (n=20) and kept in the incubator at 32±0.5 °C, 50% RH. (2) The workers were fed *ad libitum* with a sucrose solution (50% w/v in water), workers in *N. ceranae*-challenged groups were starved for 2 h one day after eclosion, and 20 workers per group were then each immobilized and fed with 5 µL of 50% sucrose solution containing 1×10^6^ spores of *N. ceranae*; these workers were isolated for 30 min in vials in the growth chamber to ensure that the sucrose solution was not transferred among individual workers and the total dosage was ingested. Workers in un-challenged groups were inoculated in an identical manner with a 50% sucrose solution without spores. Each cage was checked every 24 h and any dead workers removed. (3) Midgut tissues of *N. ceranae*-challenged and un-challenged workers were respectively selected at 7 dpi and 10 dpi, immediately frozen in liquid nitrogen and kept at −80 °C until sequencing. *N. ceranae*-challenged groups were termed as AcT1 and AcT2; un-challenged groups were termed as AcCK1 and AcCK2.

### 2.2. cDNA library construction and Nanopore long-read sequencing

Firstly, total RNA of AcCK1, AcCK2, AcT1 and AcT2 were respectively isolated using TRizol reagent (Thermo Fisher, Shanghai, China) on dry ice, and then subject to reverse transcription with Maxima H Minus Reverse Transcriptase and concentration on a AMPure XP beads. Secondly, cDNA libraries were constructed from 50ng total RNA with a 14 cycles PCR using the cDNA-PCR Sequencing Kit (SQK-PCS109) and PCR Barcoding Kit (SQK-PBK004) following the manufacturer’s protocol (Oxford Nanopore Technologies Ltd, Oxford, UK). Thirdly, the cDNA libraries were sequenced using PromethION platform (Oxford Nanopore Technologies Ltd, Oxford, UK) by Biomarker Technologies (Beijing, China).

### 2.3. Processing of long reads

Raw reads were first filtered with minimum average read quality score=7 and minimum read length=500bp; ribosomal RNA were discarded after mapping to rRNA database. Subsequently, full-length non-chemiric (FLNC) transcripts were determined by searching for primer at both ends of reads. Next, clusters of FLNC transcripts were obtained after mapping to the reference genome of *Apis cerana* (assembly ACSNU-2.0) with mimimap2 [6]; consensus isoforms were gained after polishing within each cluster by pinfish (https://github.com/nanoporetech/pinfish) and then mapped to the *A. cerana* genome (assembly ACSNU-2.0) using minimap2. Finally, mapped reads were further collapsed by cDNA_Cupcake package (https://github.com/Magdoll/cDNA_Cupcake) with min-coverage=85% and min-identity=90%; 5’ difference was not considered when collapsing redundant transcripts.

## Acknowledgments

This research was supported by the Earmarked Fund for China Agriculture Research System (No. CARS-44-KXJ7), the Science and Technology Planning Project of Fujian Province (No. 370 2018J05042), the Teaching and Scientific Research Fund of Education Department of Fujian Province (No. JAT170158), the Outstanding Scientific Research Manpower Fund of Fujian Agriculture and Forestry University (No. xjq201814), and the Scientific and Technical Innovation Fund of Fujian Agriculture and Forestry University (No. CXZX2017342, No. CXZX2017343).

